# Extracellular Electron Transfer by *Shewanella oneidensis* Controls Pd Nanoparticle Phenotype

**DOI:** 10.1101/327460

**Authors:** Christopher M. Dundas, Austin J. Graham, Dwight K. Romanovicz, Benjamin K. Keitz

## Abstract

Biological production of inorganic materials is impeded by relatively few organisms possessing genetic and metabolic linkage to material properties. The physiology of electroactive bacteria is intimately tied to inorganic transformations, which makes genetically tractable and well-studied electrogens, such as *Shewanella oneidensis*, attractive hosts for material synthesis. Notably, this species is capable of reducing a variety of transition-metal ions into functional nanoparticles, but exact mechanisms of nanoparticle biosynthesis remain ill-defined. We report two key factors of extracellular electron transfer by *S. oneidensis*, the outer membrane cytochrome, MtrC, and soluble redox shuttles (flavins), that affect Pd nanoparticle formation. Changes in the expression and availability of these electron transfer components drastically modulated particle phenotype, including particle synthesis rate, structure, and cellular localization. These relationships may serve as the basis for biologically tailoring Pd nanoparticle catalysts and could potentially be used to direct the biogenesis of other metal nanomaterials.

## Introduction

Control over cellular machinery has enabled the production of diverse compounds including pharmaceuticals, fuels, fine chemicals, and soft materials^1–3^. However, microbial engineering of metal and metal oxide products remains a significant challenge. Attributes such as electron transfer and nucleation are typically absent from organic biosyntheses, and their presence adds increased complexity to inorganic transformations. As a result, the enzymatic and metabolic factors that drive material formation in biological systems have been challenging to study and manipulate^4^. Despite the paucity of bioengineered inorganic products, microbial transformation of inorganics occurs quite frequently in nature. Several organisms generate highly functional and ordered materials, including metal nanoparticles, silicas, calcium carbonates, and metal oxides^5–7^. In these systems, coordinated protein and metabolite networks exert control over material morphology, composition, and function. For instance, the magnetotactic bacterium, *Magnetospirillum gryphiswaldense*, uses several proteins that govern nucleation, electron transfer, and vesicle formation to generate size-controlled magnetite nanoparticles^8^. This example highlights the capability of living systems to tailor inorganic structure-function relationships and suggests that exploiting naturally-occurring pathways may provide a means for designer material biosynthesis.

Electroactive bacteria are attractive hosts for inorganic materials engineering, as a diversity of soluble and insoluble inorganic substrates can be incorporated into their metabolism^9^. Whereas magnetotactic bacteria are limited to generating iron oxides, electrogens can transfer respiratory electron flux onto several metal species, including Cu(II), U(VI), Ag(I), Au(III), and Pd(II), to generate functional nanoparticles^5^. These particles have found catalytic utility in bioremediation and organic synthesis, and in some cases exhibit superior activity to those synthesized via traditional methods. However, it is generally unclear how electroactive physiology dictates the structural and functional properties of produced nanoparticles.

One electroactive bacterium, *Shewanella oneidensis* MR-1, is poised to address this issue, as it directs metabolic electron flux onto metals using a well-characterized electron transport pathway^10^. The organism’s genetic tractability has also facilitated understanding and control of this network, with knockout, complementation, and overexpression studies leading to identification of important redox-active metalloproteins and small-molecules^11^. Notably, this pathway can reduce substrates located outside the bacterial outer membrane in a process known as extracellular electron transfer (EET). Despite significant progress in applying this system towards bioremediation and microbial fuel cell engineering, elucidating the function of EET components in nanoparticle formation has proven challenging. Whereas the outer membrane cytochromes, MtrC and OmcA, are primary mediators of EET to iron oxides and electrodes^12,13^, their activity in nanoparticle formation appears largely dependent on culture conditions and the identity of metal reduced. For example, a knockout strain deficient in these proteins changed the cellular localization of produced UO_2_ nanoparticles relative to those of MR-1^14^, but had no measured effect on the kinetics of Pd or Cu nanoparticle formation^15,16^. Additional proteins have been identified as influencing nanoparticle formation rates, including hydrogenases and other periplasmic reductases^17,18^, but little is known of their influence on particle properties. Furthermore, the importance of soluble redox shuttles (flavins) in EET by *Shewanella* has become increasingly apparent^19^, but their impact on nanoparticle synthesis is wholly unexamined.

To better inform the design of material syntheses using *S. oneidensis*, we explored the role of key EET factors in the formation of Pd nanoparticles. Pd was chosen as it is one of the more well-studied nanoparticles generated by *S. oneidensis* and enabled us to draw comparisons with previous reports^15,20–22^. Pd nanoparticles are also industrially-relevant catalysts whose catalytic activity is largely dictated by their crystallographic structure and morphology^23^. Bio-based stabilization of specific Pd crystal facets was previously demonstrated via peptide-directed synthesis^24^, and we reasoned that *S. oneidensis* cells may exhibit similar control of particle nucleation and growth. In contrast to previous works, the EET machinery of *S. oneidensis* also affords the opportunity to genetically and metabolically instruct electron transfer to Pd(II) ions during nanoparticle synthesis. Thus, we tested the effect of *S. oneidensis* genotype and the role of redox-active small molecules in Pd nanoparticle formation. Here, we report two factors of *S. oneidensis* electron transport that strongly influence nanoparticle phenotype: the outer membrane cytochrome, MtrC, and soluble redox shuttles (flavins). After manipulating the bacterial concentration of these species, through control of gene expression and exogenous supplementation, respectively, we observed drastic effects on the synthesis rate, size, and cellular localization of biogenic nanoparticles. Identification of these factors provides proof-of-principle for using genetic and metabolic manipulation to tune the properties of Pd and potentially other metal nanoparticles generated by *S. oneidensis*.

## Results and Discussion

### Optimizing *S. oneidensis* Control Over Nanoparticle Biosynthesis

To drive *S. oneidensis* control over Pd nanoparticle formation, we first identified nanoparticle synthesis conditions that minimized abiotic effects and exhibited unambiguous nanoparticle phenotypes. Specifically, we examined how reaction media formulation and method of anoxic culture influenced nanoparticles formed by *S. oneidensis* MR-1. The general procedure for Pd nanoparticle biosynthesis was as follows: MR-1 was anaerobically pregrown overnight to stationary-phase in *Shewanella* Basal Medium (SBM) containing lactate/fumarate, anaerobically washed with degassed SBM, and finally used to inoculate a Pd nanoparticle reaction mixture (final OD_600_ ~0.2). Whole mount transmission electron microscopy (TEM) enabled initial assessment of different conditions, as the bacteria became palladized upon exposure to Pd(II) and adherent nanoparticles could be visualized by electron micrographs.

Choice of electron donor can greatly affect bacterial nanoparticle formation, as many metabolically accessible electron donors (e.g. formate, hydrogen) also abiotically reduce palladium ions^25^. Indeed, we found that the hydrogenous atmosphere (3%) of a humidified anaerobic glovebox caused autocatalytic reduction of palladium in the absence of bacteria. Thus, we used butyl rubber-stoppered Hungate Tubes purged with argon to maintain anaerobicity for bacterial pregrowth and nanoparticle biosynthesis reactions. As lactate is relatively inert to palladium and generates metabolic electron flux in *Shewanella*, we used it as our primary electron donor. Reaction mixtures omitting lactate or any other electron donor showed no Pd(II) reduction (Figure S1). We also assessed the influence of culture medium components that could cause abiotic reduction of Pd(II). Prior to bacterial inoculation of the reaction mixture, *S. oneidensis* was pregrown in SBM supplemented with casamino acids. When the nanoparticle reaction mixture was similarly supplemented, small Pd nanoparticles were formed on bacteria (<10 nm) (Figure S2). In contrast, mixtures omitting casamino acids had a larger nanoparticle size distribution, with particles generally falling within one of two populations: small (<10 nm) and large (~50 nm) particles. We speculated that the presence of free cysteines may cause abiotic reduction of Pd(II) and lead to fewer large particles formed on bacteria. Thus, to minimize abiotic effects we utilized reaction mixtures that lacked casamino acids.

The intracellular space is a reducing environment and leakage of promiscuous reductants (e.g. NADH, glutathione) through compromised membranes could also contribute to Pd(II) reduction. To address this, we performed viability measurements to quantify the effect of increasing concentration of Pd(II) in the *S. oneidensis* reaction mixtures. Viability was quantified using the BacLight Live/Dead stain after exposure to 10, 100, and 1000 μM Pd(II) for two hours under our standard nanoparticle synthesis conditions. *S. oneidensis* viability exhibited a dose-dependent response for the concentration range tested, varying from ~80% to ~30% viable with increasing Pd(II) concentration (Figure S3). To mitigate the effects of promiscuous reducing agents and cell death, but also balance sufficient nanoparticle yield for characterization and analysis, we chose 100 μM Pd(II) as our primary concentration for nanoparticle synthesis reactions (except where noted).

### Loss of Outer Membrane Cytochromes Decreases Pd Nanoparticle Size and Alters Cellular Localization

Pathways for both soluble and insoluble metal reduction are well-studied in *S. oneidensis*^10^. These EET pathways utilize a network of electron transfer proteins to direct metabolic electron flux from the cytoplasm, across the periplasm, and into the extracellular space (Figure 1a). As reduction of Pd(II) is necessary for nanoparticle formation, we examined the particle phenotypes of strains lacking key components of this network. The outer membrane *c*-type cytochromes, MtrC and OmcA, are critical mediators of extracellular electron transfer to metal substrates and have shown some influence on affecting the localization of produced nanoparticles^14^. We hypothesized that they may also participate in Pd(II) reduction and that a knockout strain (Δ*mtrC*Δ*omcA*) would exhibit distinct nanoparticle properties. Additionally, periplasmic hydrogenases have demonstrated Pd(II) reduction activity in *S. oneidensis* and several other bacteria^15,26,27^. Thus, we also examined the characteristics of Pd nanoparticles formed by a hydrogenase deficient strain (Δ*hydA*Δ*hyaB*).

**Figure 1.**
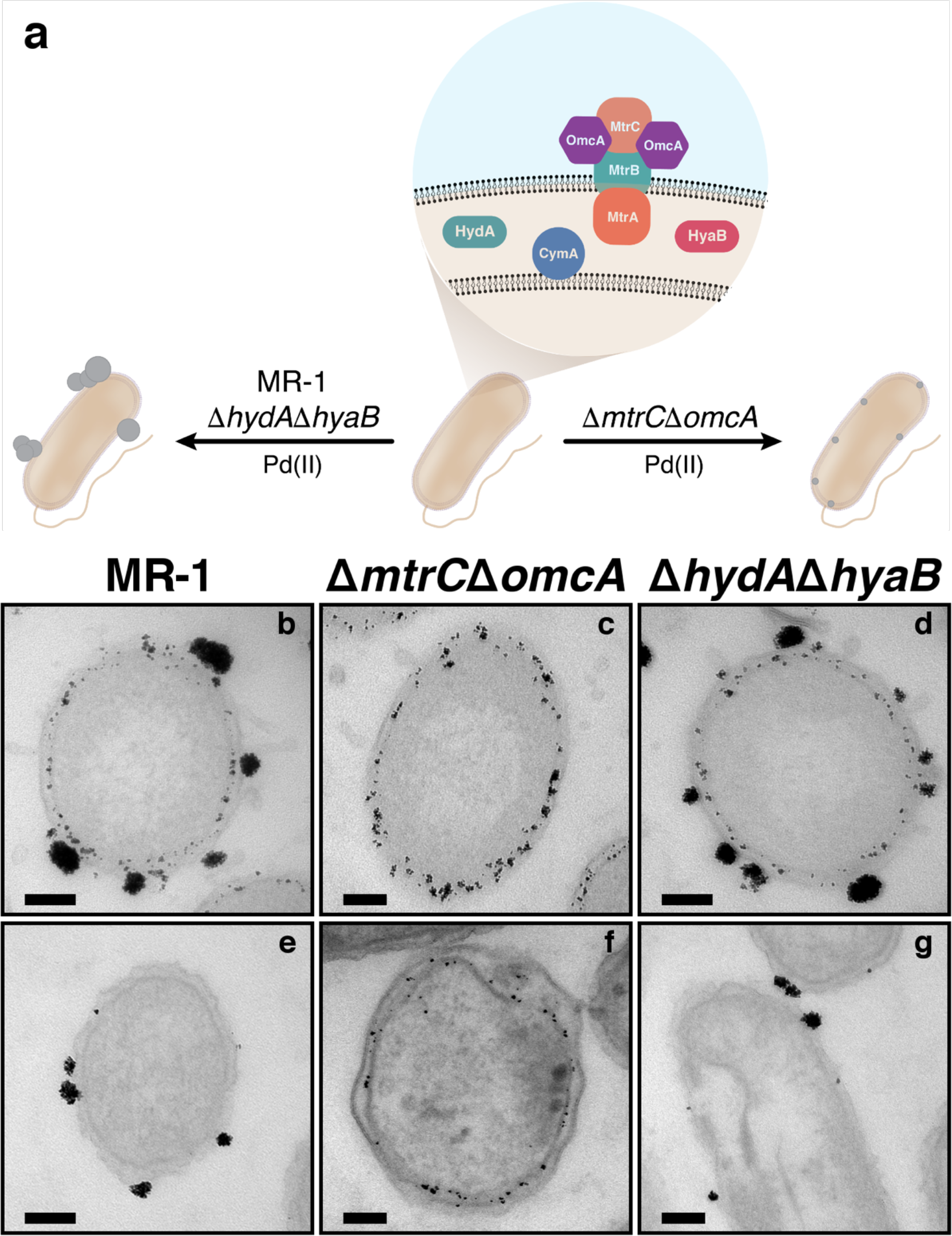
Formation of extracellular Pd nanoparticles requires outer membrane cytochromes. (a) General diagram of electron transport in *S. oneidensis* and genotypic effects on Pd nanoparticle formation. Thin section transmission electron micrographs of (b) MR-1, (c) Δ*mtrC*Δ*omcA*, and (d) Δ*hydA*Δ*hyaB* after 24 hour reactions containing 1000 μM Pd(II). (e) MR-1, (f) Δ*mtrC*Δ*omcA*, and (g) Δ*hydA*Δ*hyaB* after 2 hour reactions containing 100 μm Pd(II). Scale bars represent 100 nm.

Noticeable differences between nanoparticle reaction mixtures containing different strains were observed shortly after bacterial inoculation. Reactions inoculated with MR-1 or the Δ*hydA*Δ*hyaB* strain turned a diffuse black color that is characteristic of reduced Pd^27^, while those with the Δ*mtrC*Δ*omcA* mutant remained relatively colorless. Similarly, the cell pellets for the MR-1 and the Δ*hydA*Δ*hyaB* strains were dark black, while the Δ*mtrC*Δ*omcA* pellet appeared light grey (Figure S4). Whole mount TEM showed that these differences arose based on the types of nanoparticles generated by each strain. The large particle population observed on MR-1 was not found on the Δ*mtrC*Δ*omcA* strain and only small nanoparticles were adherent to this cytochrome double knockout (Figure S5). In contrast, nanoparticles on the Δ*hydA*Δ*hyaB* strain were identical to those on MR-1, exhibiting both large and small particle populations. As whole mount TEM cannot resolve the cellular location of particles, thin section TEM was performed. Thin sectioning of all three strains revealed that the two populations of particles not only differ in size, but also in cellular localization: smaller (<10 nm) particles localized to the periplasm and larger particle aggregates (~50 nm) adhered to the outer membrane and faced the extracellular space. Consistent with whole mount observations, MR-1 and the Δ*hydA*Δ*hyaB* strain contained both particle populations, while the Δ*mtrC*Δ*omcA* strain lacked extracellular particles (Figure 1 b-g, S6). A similar change in UO_2_ nanoparticle localization has previously been observed with this mutant^14^. Across all strains, no Pd particles were detected within the bacterial cytoplasm, indicating that reduction primarily occurred in the periplasm or extracellular space. Comparisons between bacteria palladized at 100 and 1000 μM Pd(II) showed that more particles were produced per cell at higher Pd(II) concentrations. However, nanoparticle localization was more binary at the lower Pd(II) concentration. At low Pd(II) concentrations, particles exclusively nucleated extracellularly on the outer membrane (MR-1, Δ*hydA*Δ*hyaB*) or within the periplasm (Δ*mtrC*Δ*omcA*). Together, our data suggests that low Pd(II) concentrations accentuate genotypic differences, whereas decreased cell viability at high Pd(II) concentrations (Figure S3) results in more promiscuous Pd(II) reduction.

In addition to TEM, powder x-ray diffraction (PXRD) was used to characterize the properties of bacterial Pd nanoparticles. PXRD patterns can reveal the chemical identity and relative size of analyzed nanomaterials. Differences between *S. oneidensis* strains were detected by PXRD, and corroborated trends observed by TEM (Figure S7). While nanoparticles formed by MR-1 and the Δ*hydA*Δ*hyaB* mutant had comparable patterns, broader peaks were observed for those from the Δ*mtrC*Δ*omcA* strain, indicating smaller grain size of Pd nanocrystallites as predicted by the Scherrer equation^28^. The observed peak locations were consistent with previous Pd-*Shewanella* studies and with abiotic Pd(0) patterns^20^, verifying that peaks were a result of palladization. These results further demonstrate the importance of outer membrane cytochromes in controlling nanoparticle properties and show that PXRD can be used to distinguish genotypic differences.

The existence of the two particle populations illuminates how bacterial membranes and outer membrane cytochromes influence nanoparticle nucleation and growth. Cellular compartmentalization appears to physically constrain nanoparticle structure, as evidenced by intracellular particles limited in size by periplasmic dimensions (<50 nm), while extracellular particles further aggregated (>50 nm). Importantly, bacterial genotype controlled particle localization, since loss of outer membrane cytochromes MtrC and OmcA was coincident with loss of outer membrane particles (Figure 1c, 1f). Nanoparticle reaction mixtures that replaced *S. oneidensis* with a cytochrome-lacking bacterium, *Escherichia coli* MG1655, similarly had no large extracellular particles formed on the bacteria (Figure S8). Additionally, post-palladization viability studies performed on MR-1 and the Δ*mtrC*Δ*omcA* strain showed that changes in cell viability do not account for the differences between particle phenotypes (Figure S9). Together, this indicates that the outer membrane cytochromes are critical for extracellular Pd nucleation and likely serve as sites for electron transfer to Pd(II).

### Loss of Outer Membrane Cytochromes Attenuates Pd(II) Reduction

Generally, the structures of biosynthesized metabolites are invariant to the rate at which they are produced. In contrast, metal ion reduction rates can exert significant influence over the size and morphology of produced nanoparticles^29^. To assess whether reduction kinetics could explain differences in Pd particle size/localization between *S. oneidensis* strains, we quantified Pd(II) concentrations over the course of nanoparticle synthesis.

With all strains, extracellular Pd(II) concentrations were measured as ~50 μM at the first time-point (starting concentration was 100 μM), indicating rapid Pd(II) adsorption to cells upon bacterial inoculation (Figure 2). Reaction mixtures with *E. coli* MG1655 also showed Pd(II) adsorption to cells, but no Pd(II) reduction was detected on the same time scale (Figure S10). Following the initial adsorption event, we measured a comparable first-order Pd(II) reduction rate between MR-1 and the hydrogenase deficient mutant when lactate was utilized as an electron donor (Figure 2, S11). Although kinetic differences between these strains have previously been measured when utilizing formate^15^, less metabolic electron flux is directed to the quinone pool from formate catabolism^30^. This suggests that the outer membrane cytochromes are primarily responsible for Pd(II) reduction when electron flux can be directed through the Mtr pathway, which utilizes the quinone pool. Accordingly, the Δ*mtrC*Δ*omcA* strain showed significantly attenuated Pd(II) reduction relative to MR-1. These results differed from a previous study, where a strain lacking these cytochromes had a rate of reduction rate that was identical to MR-1^15^. We attribute these conflicting observations to the significant effects that buffering capacity can exert on the kinetics of Pd(II) reduction by *S. oneidensis*. We note that our bacterial pregrowth and nanoparticle reaction mixtures utilized substantially more buffered media (100 mM HEPES) than this previous study (30 mM HEPES). Even under our conditions, Pd(II) reduction rate greatly decreased with increasing age of dissolved HEPES in pregrowth and reaction media (Figure S12). Cell growth can alter culture medium acidity, and the pH stability of strongly buffered media may counter physiological shifts such as cytochrome downregulation or inactivity that otherwise occurs^31^. Alternatively, the loss of MtrC and OmcA increased the variability between biological replicates, which could arise from the increased role of promiscuous reductants as opposed to specific electron transport pathways. This may have further obscured kinetic differences between strains in previous studies. Taken together, our results are consistent with previously purported nanoparticle formation mechanisms by *S. oneidensis*: Pd(II) adsorbs to cells, Pd(II) is reduced, and Pd nanoparticles nucleate^15^. Notably, our results highlight the importance of outer membrane cytochromes in the latter two steps.

**Figure 2.**
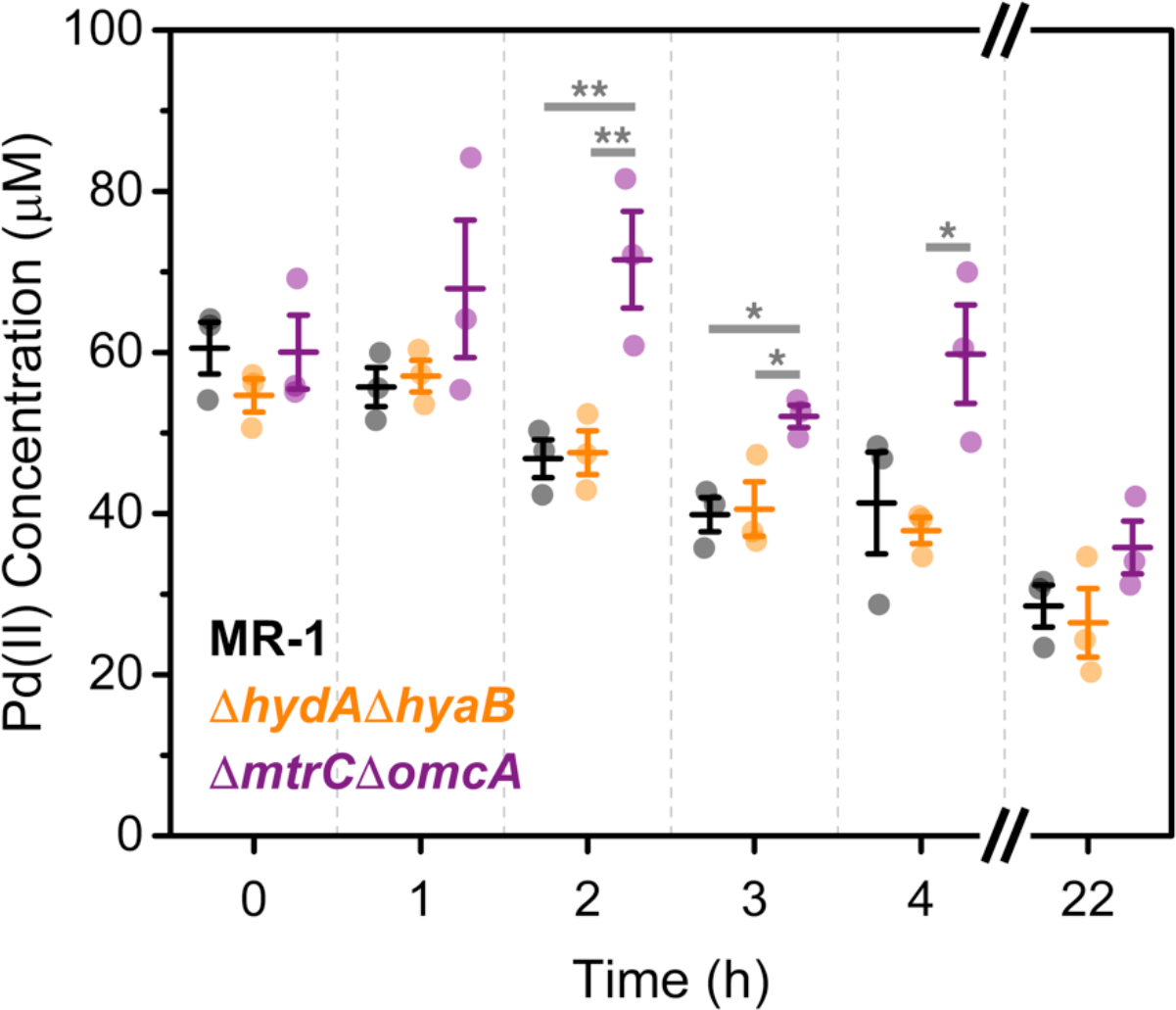
Loss of outer membrane cytochromes attenuates Pd(II) reduction rate. Extracellular Pd(II) concentrations during nanoparticle synthesis for MR-1, Δ*mtrC*Δ*omcA*, and *ΔhydAΔhyaB*. Raw data is overlaid on bars representing the mean ± S.E. * p < 0.05, ** p < 0.01, n = 3.

### MtrC Expression Controls Extracellular Pd Nanoparticle Formation

To better understand the role of MtrC and OmcA in Pd(II) reduction and nanoparticle formation, we complemented the Δ*mtrC*Δ*omcA* strain with one of three plasmids that was either an empty control vector or vectors carrying *mtrC* and *omcA* under constitutive control^32^.

We first tested whether the presence of either MtrC or OmcA was sufficient to restore the extracellular particle phenotype observed with MR-1. Whole mount TEM showed that Δ*mtrC*Δ*omcA* strains carrying either an empty vector or an *omcA*-expressing vector were unable to form large nanoparticles that are characteristic of extracellular localization (Figure S13). However, large particles were observed on the double knockout strain complemented with a plasmid carrying *mtrC*. Thin section TEM confirmed that large particles formed by this *mtrC*-complemented strain were extracellular and adherent to the bacterial outer membrane (Figure 3, S14). MR-1 carrying an empty plasmid exhibited an identical phenotype, which validated that MtrC expression is responsible for extracellular particle nucleation and plasmid maintenance does not interfere with nanoparticle formation. These results are consistent with previous reports that describe MtrC functioning in the Mtr pathway^12^. In these studies, MtrC expression restored EET to Fe(III) species in outer membrane cytochrome deficient strains, while OmcA only partially contributed to EET. OmcA may act as a non-critical helper protein that requires MtrC to receive electrons for subsequent metal reduction. Our results suggest that this mechanism is also true for Pd(II) reduction.

**Figure 3.**
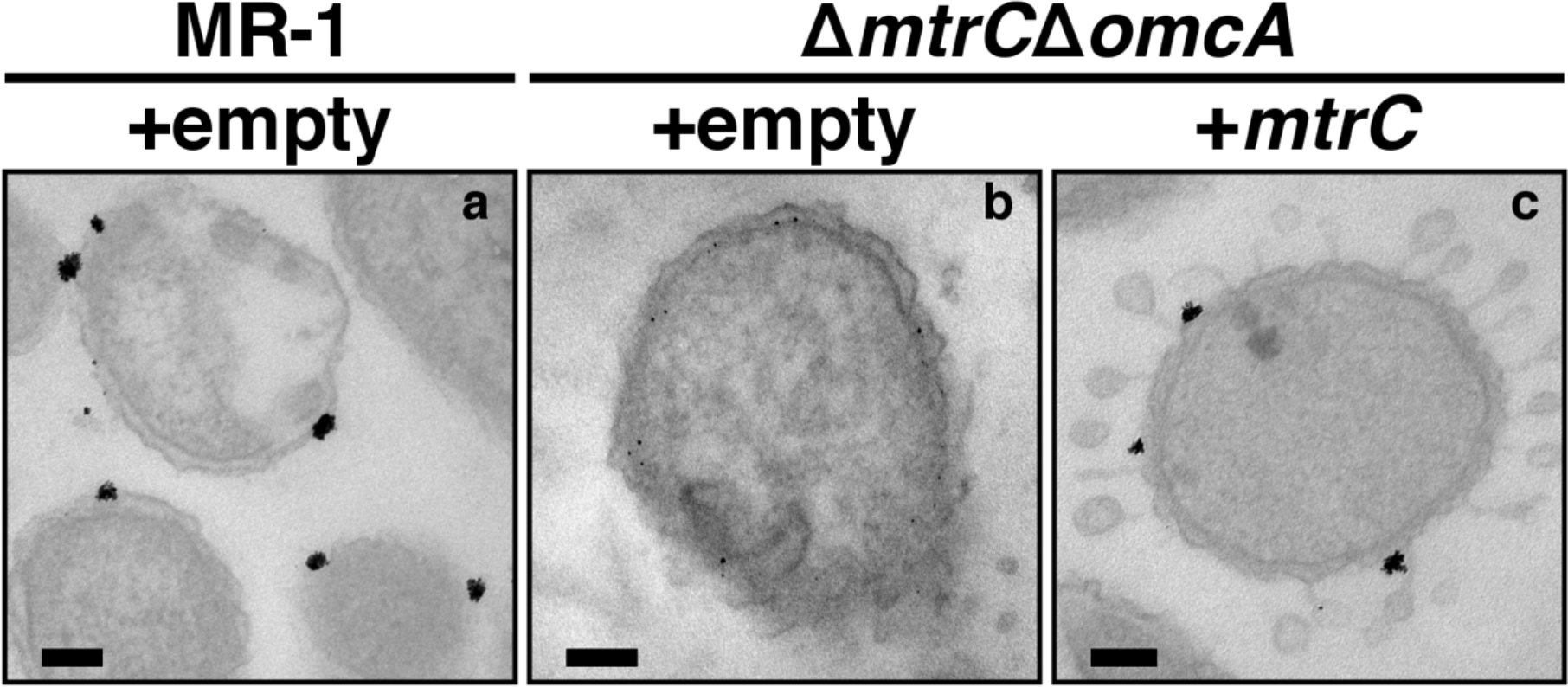
MtrC expression controls extracellular localization of Pd nanoparticles. Thin section transmission electron micrographs of (a) MR-1 with an empty vector and Δ*mtrC*Δ*omcA* with an (b) empty vector or (c) *mtrC* vector after 2 hours with 100 μm Pd(II). Scale bars represent 100 nm.

We next measured Pd(II) reduction kinetics with the *mtrC*-complemented mutant to assess whether wild-type Pd(II) reduction was rescued. Indeed, reduction was substantially faster with the *mtrC*-complemented mutant relative to the same mutant carrying an empty plasmid (Figure 4, S11). Moreover, the *mtrC*-complemented mutant exhibited a faster Pd(II) reduction rate compared to MR-1 carrying an empty plasmid. This is likely due to different MtrC expression levels between our engineered strain and MR-1. Taken together with our microscopy and knockout results, our complementation studies confirm that MtrC is a critical enabler of Pd(II) reduction and extracellular nanoparticle nucleation. The complementation kinetics also suggest that manipulating expression levels of MtrC may be a viable means for tuning Pd(II) reduction rates or controlling Pd nanoparticle localization.

**Figure 4.**
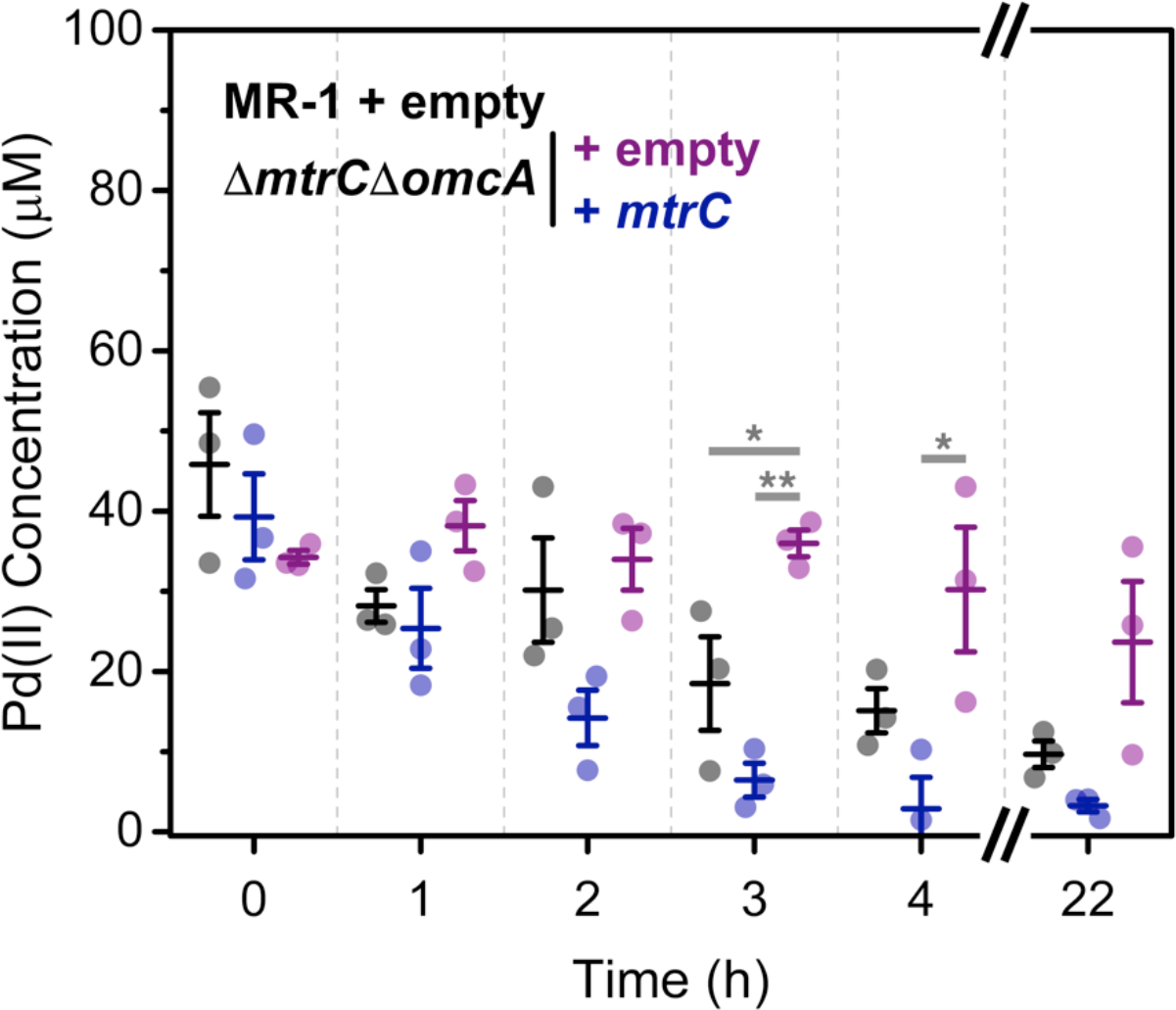
MtrC expression rescues Pd(II) reduction rate. Extracellular Pd(II) concentrations during nanoparticle synthesis for plasmid-complemented *S. oneidensis* strains. Raw data is overlaid on bars representing the mean ± S.E. * p < 0.05, ** p < 0.01, n = 3.

### Flavins Decrease Outer Membrane-Nanoparticle Size

Flavins are a family of redox-active small-molecules that function in *E. coli, Shewanella*, and other microorganisms^33^. In *Shewanella*, they have been implicated in a unique role of accelerating electron transfer to insoluble metal substrates^19^. Strains lacking a key periplasmic flavin exporter (Δ*bfe*) exhibit significantly attenuated electron transfer to iron oxides and electrodes^34^. It has been proposed that flavin binding to MtrC and other outer membrane cytochromes is critical for enabling direct-contact electron transfer to insoluble substrates^35^. Furthermore, flavins are known chelators that can bind to a variety of soluble and insoluble metal species^19,36^. These findings suggest that flavin concentration may also play a role in nanoparticle synthesis by influencing electron transfer and/or instructing metal binding.

We measured Pd nanoparticle properties in response to flavin supplementation of the nanoparticle reaction mixture. Nanoparticle size decreased in a dose-dependent manner with increasing riboflavin concentration (1, 10, 100 μM), when analyzed by whole mount TEM (Figure S15). Similarly, supplementing reaction mixtures with flavin mononucleotide (100 μM) caused a small-particle phenotype (Figure S16). Thin section TEM demonstrated that in the presence of exogenous flavins, Pd nanoparticles were localized to both the periplasm and outer membrane. However, extracellular particles were significantly smaller (<10 nm) and in greater number per cell compared to the non-supplemented reaction mixtures (Figure 5). This differs substantially from strains lacking *mtrC*, which generated smaller particles exclusively confined to the periplasm. Riboflavin and flavin mononucleotide are thought to directly mediate electron transfer by binding to OmcA and MtrC^35^, respectively, and an increase in cytochrome-binding may explain the greater number of Pd nanoparticles on the outer membrane. In contrast to flavins, supplementing other respiration substrates (sodium fumarate and Fe(III)-citrate) showed no effect on nanoparticle size (Figure S17, S18). Additionally, we measured no effect on cell viability when supplementing with flavins, which discounts the influence of stress response on affecting nanoparticle formation (Figure S19). These results support our hypothesis that, unlike other metabolites, flavins play a role in either electron transfer to Pd(II), particle nucleation/growth, or both.

**Figure 5.**
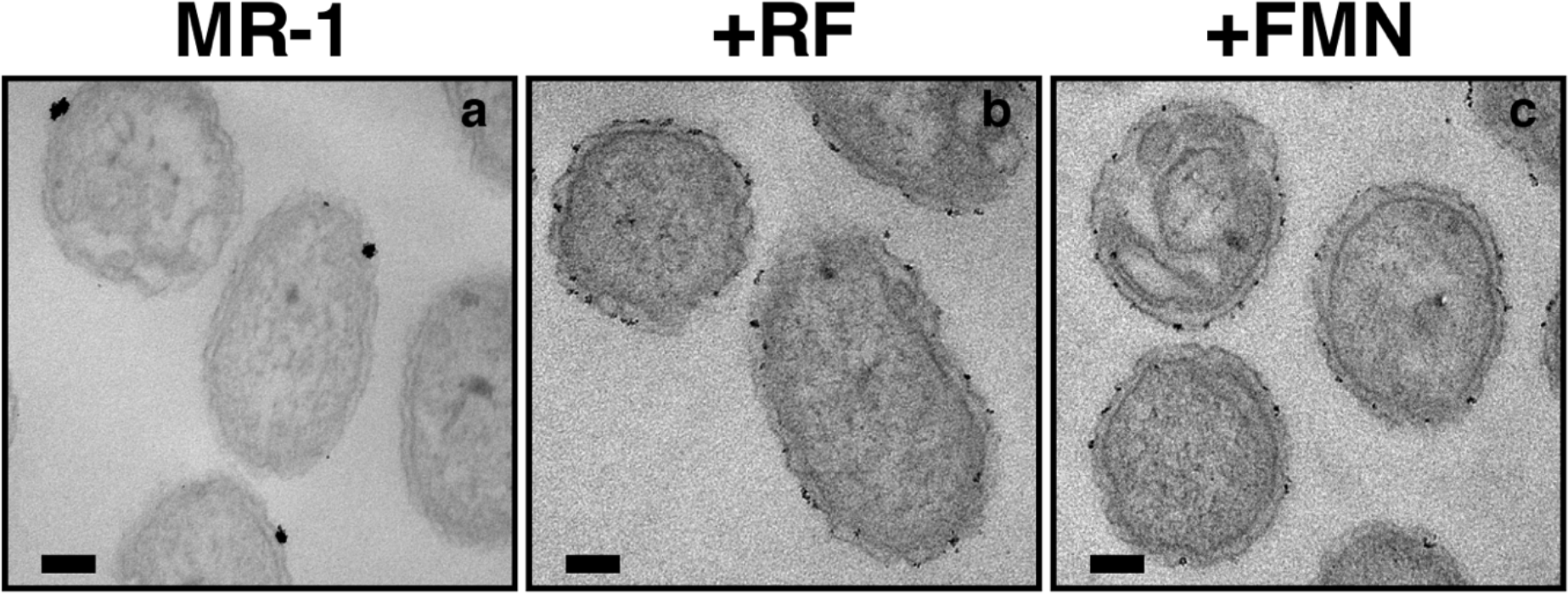
Flavin supplementation decreases size of extracellular Pd nanoparticles. Thin section transmission electron micrographs of (a) MR-1 and MR-1 with (b) 100 μm riboflavin (RF) or (c) 100 μm flavin mononucleotide (FMN) after 2 hours with 100 μM Pd(II). Scale bars represent 100 nm.

To further investigate this mechanism, we measured Pd(II) reduction kinetics in the presence of 100 μM of exogenous riboflavin or flavin mononucleotide. Pd(II) reduction rates of MR-1 supplemented with riboflavin were slightly, yet significantly, slower compared to samples omitting supplementation. (Figure 6). While previous findings suggest that flavins are not critical mediators of electron transfer to soluble metal acceptors^32,34^, their function in precipitation of soluble Pd appears more nuanced and with competing factors at play. Attenuated Pd(II) reduction could be explained by flavins binding and sequestering soluble Pd(II). Flavins could also act as capping agents that passivate growing nanoparticle surfaces and hinder further Pd(II) reduction^37^. Indeed, the presence of riboflavin or flavin mononucleotide in an abiotic synthesis of Pd nanoparticles using NaBH_4_ prevented significant particle aggregation relative to a flavin-free control (Figure S20). Overall, our results demonstrate that flavin concentration exerts control over Pd nanoparticle size and reduction rate without affecting their cellular localization.

**Figure 6.**
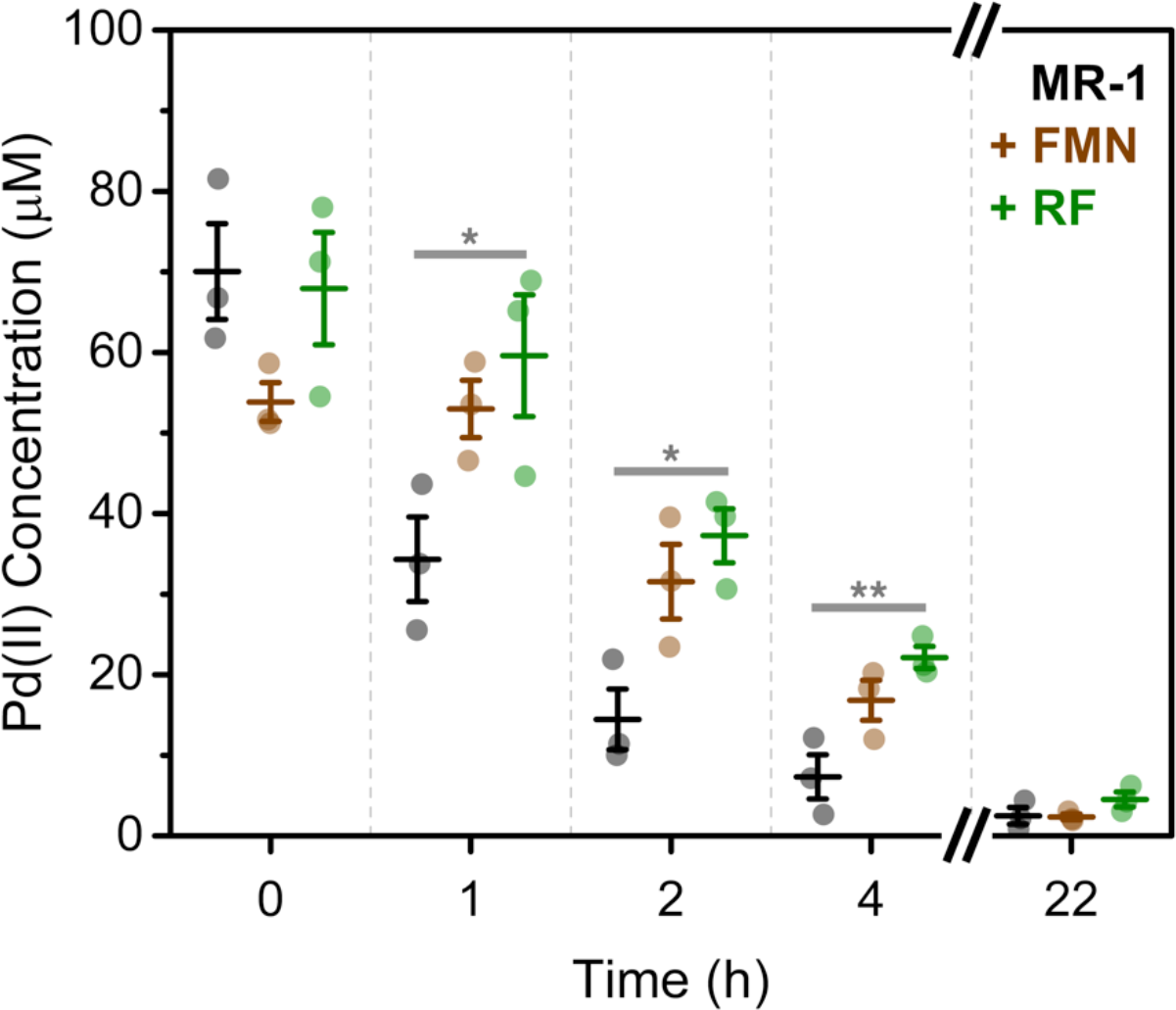
Flavin supplementation attenuates Pd(II) reduction rate by *S. oneidensis*. Extracellular Pd(II) concentration for unsupplemented MR-1 and MR-1 supplemented with 100 μM riboflavin (RF) or 100 μm flavin mononucleotide (FMN). Raw data is overlaid on bars representing the mean ± S.E. * p < 0.05, ** p < 0.01, n = 3.

## Implications

To summarize, we determined that the EET components, MtrC and flavins, can exert significant influence over Pd nanoparticle formation by *Shewanella oneidensis*. By adjusting their concentration in the bacterial environment, significant changes were actuated in Pd nanoparticle synthesis rate, size, and cellular localization.

Knockout and complementation experiments confirmed that the outer membrane cytochrome, MtrC, is a critical enabler of rapid Pd(II) reduction and extracellular nanoparticle nucleation. Notably, expression of MtrC via an engineered plasmid led to Pd(II) reduction rates that were faster than wild-type controls. Given this observation, coupled with the loss of extracellular particles on *mtrC* knockouts, we hypothesize that more fine-tuned expression of MtrC may correlate to different numbers and sizes of produced nanoparticles. This strategy may also be adapted to standard chassis organisms that express functional MtrC, such as engineered *E. coli* strains^38^, or other electroactive bacteria possessing Pd(II)-reducing outer membrane cytochromes (e.g. *Geobacter sulfurreducens*^39,40^. Alternatively, nanoparticle formation could be varied by directly engineering the MtrC polypeptide sequence, either through rational or evolutionary mutagenesis. Altering the redox-potential and catalytic activity of heme-containing proteins is well-precedented^41^, and key mutations could be applied to identify/tune the Pd(II) reduction activity of MtrC hemes^42^. Additionally, a binding-site for iron oxide nanoparticles to the outer membrane cytochrome, MtrF, has recently been elucidated using protease footprinting^43^, and similar methods could be used to identify and affect residues involved in particle nucleation by MtrC.

We observed a decrease in the size of Pd nanoparticles with increasing concentration of supplemented flavins to nanoparticle reaction mixtures. Reduction kinetics again appear to play a role in nanoparticles produced in the presence of flavins; however, unlike the effect of *mtrC* deletion, nanoparticles formed with flavins were extracellularly bound to the bacterial membrane. Given our understanding of flavin interaction with soluble metals and metal surfaces, it is likely that increased flavin concentration alters nanoparticle dimensions through binding and nucleation interactions. *Shewanella*-secreted flavins have been measured as ~0.1-1.5 μM in culture supernatants^34,44^, therefore our supplementations (100 μM) far exceeded physiologically generated concentrations. However, strains with additional flavin biosynthesis pathways have increased secreted concentrations to as high as ~25 μM^45^. Further pathway engineering would likely increase this concentration and potentially allow for stimuli-responsive pathways to control nanoparticle size. Flavin binding to outer membrane cytochromes likely also plays a role in the observed particle phenotypes^35^, and we speculate that modulating flavin-cytochrome binding affinities through protein engineering could produce similar effects without exogenous supplementation^42^. Overall, our results shed further light on the role of flavins in *S. oneidensis*-metal chemistry and demonstrate another means of tunability for biological nanoparticle synthesis.

Biological generation of inorganic structures most commonly arises in nature through the actions of complex protein and metabolic networks. However, engineered systems have been described where relatively simple biological interactions (e.g. amyloid fibers^3^, phage scaffolds^46^) can direct nanomaterial morphologies. We present an alternative strategy that uses single genetic or metabolic factors of viable *S. oneidensis* cells to control Pd nanoparticle phenotype. Our results, combined with the genetic tractability of *S. oneidensis*, suggest that it may serve as an intriguing model platform for studying nanoparticle biogenesis and engineering these particles for catalysis and sensing applications.

## Materials and Methods

### Chemicals and Reagents

Sodium tetrachloropalladate(II) trihydrate (Na_2_PdCl_4_·3H_2_O, Strem Chemicals), argon (Ar, Airgas), 4-(2-Pyridylazo)resorcinol (C_11_H_9_N_3_O_2_, Alfa Aesar), HEPES Free Acid Powder (C_8_H_18_N_2_O_4_S, Sigma-Aldrich), potassium phosphate dibasic (K_2_HPO_4_, Sigma-Aldrich), potassium phosphate monobasic (KH_2_PO_4_, VWR), sodium chloride (NaCl, VWR), ammonium sulfate (NH_4_)_2_SO_4_, Fisher Scientific), magnesium(II) sulfate heptahydrate (MgSO_4_·7H_2_O, Sigma-Aldrich), EDTA acid disodium salt dihydrate (C_10_H_14_N_2_Na_2_O_8_·2H_2_O, VWR), manganese(II) sulfate monohydrate (MnSO_4_·H_2_O, VWR), ferrous sulfate heptahydrate (FeSO_4_·7H_2_O, Alfa Aesar), cobalt(II) nitrate hexahydrate (Co(NO_3_)_2_·6H_2_O, Strem Chemicals), calcium chloride dihydrate (CaCl_2_·2H_2_O, Sigma-Aldrich), zinc(II) sulfate monohydrate (ZnSO_4_·H_2_O, Strem Chemicals), cupric sulfate pentahydrate (CuSO_4_·5H_2_O, VWR), aluminum potassium sulfate (AlK(SO_4_)_2_, Acros Organics), boric acid (H_3_BO_3_, VWR), sodium molybdate dihydrate (Na_2_MoO_4_·2H_2_O, Beantown Chemical), sodium selenite (Na_2_SeO_3_, Acros Organics), sodium tungstate dihydrate (Na_2_WO_4_·2H_2_O, Alfa Aesar), nickel(II) chloride hexahydrate (NiCl_2_·6H_2_O, Alfa Aesar), D-(+)-glucose (C_6_H_12_O_6_, Sigma-Aldrich), flavin adenine dinucleotide disodium salt hydrate (C_27_H_31_N_9_Na_2_O_15_P_2_·xH_2_O, Sigma-Aldrich), riboflavin (C_17_H_20_N_4_O_6_, Alfa Aesar), kanamycin sulfate (C_18_H_38_N_4_O_15_S, Growcells) and casamino acids (VWR) were used as received. For thin section preparation, cacodylate buffer, glutaraldehyde (50%), paraformaldehyde (16%), potassium ferrocyanide, and osmium tetroxide, and epoxy resin solutions were obtained from Electron Microscopy Sciences. Sodium DL-lactate (C_3_H_5_NaO_3_, VWR, 60% in water) was filtered using 0.2 μm PES filters. Sodium fumarate (Na_2_C_4_H_2_O_4_, VWR) was diluted in H_2_O and then filtered using 0.2 μm PES filters. LB Lennox agar powder (VWR) was dissolved in H_2_O and sterilized at 121°C and 100 kPa for 1 h. Kanamycin sulfate stock was dissolved in H_2_O and filter sterilized using 0.2 μm PES filters. When preparing *Shewanella* Basal Medium, all components were added to H_2_O and filter-sterilized using 0.2 μm PES filters.

### Bacterial Strains and Culture

Bacterial strains, plasmids, and their sources are listed in Table **S1**. Except where noted, liquid cultures and nanoparticle biosynthesis reactions occurred in autoclaved Hungate culture tubes capped with a butyl-rubber stopper and plastic screw-cap. Pregrowth for nanoparticle biosynthesis reactions was prepared as follows. Bacterial stocks stored in 20% glycerol at −80 °C were freshly streaked onto LB agar plates (containing 25 μg mL^−1^ kanamycin when appropriate) and incubated aerobically for ~18 hours at 30 °C (*S. oneidensis*) or 37 °C (*E. coli*). Colonies were picked from these plates and used to inoculate Hungate tubes containing pregrowth medium that was comprised of *Shewanella* Basal Medium with 100 mM HEPES^47^, 5 mL L^−1^ trace metal mix^48^, 0.5% casamino acids, 20 mM sodium lactate (*S. oneidensis*) or 20 mM glucose (*E. coli*), and 40 mM sodium fumarate. After inoculation, tubes were incubated at 30 °C and shaken for ~18 hours at 250 rpm. Anoxic conditions were maintained by bubbling all solutions, flushing tube headspace, and overpressuring Hungate tubes (>15 psi) with argon. During transient exposure to atmosphere (during inoculation and between washes/centrifugations), solutions were strictly handled under a gassing cannula that jettisoned argon.

### Nanoparticle Biosynthesis

To prepare 3 mL of a standard nanoparticle biosynthesis reaction, 4 mL of anaerobic pregrowth was started the previous day. Once pregrowth reached stationary-phase, the entire volume was transferred to a 15 mL conical tube and centrifuged at 3400xg for 1 h. The supernatant was removed, 1 mL of degassed SBM containing trace metal mix was used to resuspend the pellet, and this suspension was subsequently transferred to a sterile 1.5 mL microcentrifuge tube for centrifugation at 6000xg for 20 minutes. A second 1 mL SBM wash/centrifugation was performed and the bacterial suspension was concentrated to an OD_600_ of 3.0 in 200 μL, which was to be later used as reaction inoculum. 2.8 mL of reaction volume was prepared by mixing 8.55 μL of sodium lactate (60% w/w) and 30 μL of freshly prepared 10 mM Na_2_PdCl_4_•3H_2_O (dissolved in autoclaved Milli-Q H_2_O) to 2.76 mL of SBM containing trace metals. This mixture was bubbled with argon and then transferred to sealed Hungate tubes. Prior to inoculation, tube headspace was flushed with additional argon for 15 minutes. Using sterile syringe needles, the entire bacterial inoculum volume was injected into to the reaction mixture, setting the final OD_600_ to 0.2., and the first time point was simultaneously aliquoted. The headspace of reaction mixtures was briefly flushed with argon for 1 minute, and incubated inside a shaker at 30 °C and 250 rpm. All time point aliquots (~300 μL) were flash frozen in liquid N_2_ and store at -20 °C for later analysis. Reactions with higher concentrations of Pd(II) or 0.5% casamino acids were performed when indicated. For reactions inoculated with plasmid-complemented knockouts, kanamycin was supplemented to mixtures at a concentration of 25 μg mL^−1^. For *E. coli*-inoculated reactions, sodium lactate was replaced with 20 mM glucose and reactions were incubated at 37 °C. As noted in the Results and Discussion, the age of HEPES in SBM pregrowth and reaction mixtures significantly affected Pd(II) reduction. Therefore, HEPES was dissolved in SBM for pregrowth and nanoparticle reaction mixtures and used within 2 weeks.

### Transmission Electron Microscopy

Palladized bacteria were prepared as described above. Nanoparticle synthesis was stopped after 2 hours (100 μM Pd(II)) or 24 hours (1 mM Pd(II)), and cells were centrifuged/washed twice. In early experiments, cells were washed with SBM containing trace metals as to minimize osmotic shock, but we later found that autoclaved H_2_O could be utilized with minimal change in cell shape and improved electron micrograph contrast. For whole mount TEM, washed cells were concentrated to an OD_600_ of ~2.7 and 5 μL was drop-cast onto glow-discharged 200 mesh carbon-coated copper TEM grids (Electron Microscopy Sciences). For thin section TEM, cells were fixed prior to sectioning and imaging. First, a 100 mL cation stock solution was prepared that contained CaCl_2_·2H_2_O (0.3 g) and MgSO_4_·7H_2_O (1 g). Next, aldehyde fixation solution was prepared by mixing 1 mL of cation stock with 5 mL of cacodylate buffer (0.2 M, pH 7.4), 0.8 mL of glutaraldehyde (50%), 1.25 mL of paraformaldehyde (16%), and 2 mL of H_2_O. The washed cell pellet was resuspended in 1 mL of aldehyde fixation solution and incubated at room temperature for 4 hours. After, cells were washed three times in 0.1 M cacodylate buffer. An osmium tetroxide fixation solution was prepared immediately before use by mixing in a 1:1 ratio of 4% potassium ferrocyanide in 0.2 M cacodylate buffer (also prepared immediately before use) and 4% osmium tetroxide. The cell pellet was resuspended in 1 mL osmium textroxide fixation solution and incubated for 4 hours on ice. Next, the pellet was resuspended in H_2_O for 10 minutes, spun-down, and repeated until the supernatant was clear (~10 washes required). Subsequently, the pellet was dehydrated in 50, 70, 95%, and 100% (twice) ethanol, for 15 minutes at each dehydration. After the final 100% ethanol wash, the pellet was washed in acetone for 15 minutes, twice. Next, the pellet was infiltrated with 30% epoxy resin in 100% acetone, 66% resin in 100% acetone, and finally 100% resin (twice), for ~16 hours with each infiltration. Finally, the pellet was placed in fresh epoxy resin, cast into a sectioning-block mold, and heated at 65 °C for 2 days. Sectioning was accomplished using a Leica UltraCut Microtome and sections were placed on copper TEM grids to for imaging. Whole mount and thin section TEM was performed using a FEI Tecnai Transmission Electron Microscope.

### Pd(II) Quantification

Extracellular Pd(II) concentrations were primarily quantified using the colorimetric 4-(2-pyridylazo)-resorcinol (PAR) assay^49^, the general procedure is as follows. A 0.1% PAR stock solution was freshly prepared by dissolving 25 mg PAR in 1 mL 1% NaOH, and then adding this solution to 24 mL MilliQ H_2_O. A stock solution of 100 mM EDTA was also prepared and the pH was adjusted to 10.5 using NaOH. Frozen nanoparticle reaction mixture aliquots (~300 μL) were quickly thawed and centrifuged at 10,000xg for 5 minutes to pellet bacteria. Next, 100 μL of each reaction mixture supernatant was added to microcentrifuge tubes containing 500 μL of the 100 mM EDTA stock and 5 μL of the 0.1% PAR stock. This solution was heated at 80 °C for 10 minutes, cooled at 4 °C for 20 minutes, and absorbance at 515 nm was measured using a CLARIOStar platereader (BMG Labtech). Standard curves were prepared in SBM containing trace metals over a range of 0-125 μM Pd(II). Flavins exhibit absorbance overlapping that of PAR, which obscured Pd(II) quantification using this method. Therefore, inductively coupled plasma mass spectrometry (ICP-MS) was used to obtain reduction kinetics in the presence of flavins. To prepare samples for ICP-MS, aliquots were thawed, centrifuged at 10,000xg for 5 minutes, and the supernatant mixed in a 1:1 ratio with strong (~70%) nitric acid. Samples were further diluted 80-fold in weak (~2%) nitric acid to bring total dissolved solids to ppm levels and then analyzed using an Agilent 7500ce ICP-MS. Pd(II) concentration at each time point was then calculated based on the degree of sample dilution.

### Powder X-Ray Diffraction

To analyze biogenic particles by PXRD, 10 mL of nanoparticle reaction mixtures using 1000 μM Pd(II) were prepared and inoculated with MR-1, Δ*mtrC*Δ*omcA*, or Δ*hydA*Δ*hyaB*. After ~18 hours, cells were washed by centrifuging at 3400xg for 40 minutes, removing supernatant, and adding equal volume of autoclaved MilliQ H_2_O. Two more washes were performed, and after the supernatant was removed in the final wash, 1 mL of acetone was used to resuspend the pellet. This acetone-cell suspension was transferred to a microcentrifuge tube and centrifuged for 20 minutes at 6000xg. Acetone was again used to resuspend cells, and this suspension was vacufuged to aspirate the supernatant and pellet dried cell mass. X-ray diffraction (XRD) was performed on a Rigaku R-Axis Spider diffractometer with an image plate detector using Cu Kα radiation (λ=1.54 Å) and a graphite monochromator. XRD samples were prepared by mixing a small amount of dried cell mass with a droplet of mineral oil followed by mounting on a cryoloop.

### Statistical Analysis

Unless otherwise noted, data are reported as mean ± S.E. of n = 3 replicates, as this sample size was sufficiently large to detect significant differences in means. Significance was calculated using a two-tailed unpaired Student’s t-test (α=0.05) and OriginPro Software (OriginLab, Northhampton, MA).

## Author Contributions

C.M.D. and B.K.K. conceived the project; C.M.D. and A.J.G. performed the experiments. D.K.R. assisted with electron microscopy preparation and experiments. C.M.D., A.J.G., and B.K.K. analyzed the results. C.M.D., A.J.G., and B.K.K. wrote the manuscript.

## Acknowledgements

This work was supported by the Welch Foundation (Grant No. F-1929). We would like to thank Prof. Jeffrey Gralnick (U. Minnesota) for generously providing *S. oneidensis* strains Δ*mtrC*Δ*omcA* and Δ*hydA*Δ*hyaB* and Prof. Lydia Contreras (U. Texas at Austin) for generously providing *E. coli* MG1655. We would like to acknowledge ICMB Core Facilities at the UT Austin for use of their Tecnai Transmission Electron Microscope and performing DNA sequencing. We also acknowledge Dr. Nathaniel R. Miller and the Jackson School of Geosciences at UT Austin for support with ICP-MS experiments.

## Notes

The authors declare no competing financial interests.

